# Conditioned fear reactions are associated with gray matter density but not cortical microstructure

**DOI:** 10.1101/2020.12.18.423236

**Authors:** Christoph Fraenz, Dorothea Metzen, Christian J. Merz, Helene Selpien, Nikolai Axmacher, Erhan Genç

**Affiliations:** Department of Biopsychology, Institute of Cognitive Neuroscience, Faculty of Psychology, Ruhr University Bochum, 44801 Bochum, Germany; Department of Cognitive Psychology, Institute of Cognitive Neuroscience, Faculty of Psychology, Ruhr University Bochum, 44801 Bochum, Germany; Department of Neuropsychology, Institute of Cognitive Neuroscience, Faculty of Psychology, Ruhr University Bochum, 44801 Bochum, Germany; Department of Psychology and Neurosciences, Leibniz Research Centre for Working Environment and Human Factors (IfADo), 44139 Dortmund, Germany

## Abstract

Research has shown that fear acquisition, in reaction to potentially harmful stimuli or situations, is characterized by pronounced interindividual differences. It is likely that such differences are evoked by variability in the macro- and microstructural properties of brain regions involved in the processing of threat or safety signals from the environment. Indeed, previous studies have shown that the strength of conditioned fear reactions is associated with the cortical thickness or volume of various brain regions. However, respective studies were exclusively targeted at single brain regions instead of whole brain networks. Here, we tested 60 young and healthy individuals in a differential fear conditioning paradigm while they underwent fMRI scanning. In addition, we acquired T1-weighted and multi-shell diffusion-weighted images prior to testing. We used task-based fMRI data to define global brain networks which exhibited increased BOLD responses towards CS+ or CS- presentations, respectively. From these networks, we obtained mean values of gray matter density, neurite density, and neurite orientation dispersion. We found that mean gray matter density averaged across the CS+ network was significantly correlated with the strength of conditioned fear reactions quantified via skin conductance response. Measures of neurite architecture were not associated with conditioned fear reaction in any of the two networks. Our results extend previous findings on the relationship between brain morphometry and fear learning. Most importantly, our study is the first to introduce neurite imaging to fear learning research and discusses how its implementation can be improved in future research.

## 1. Introduction

In our everyday lives, we are constantly confronted with threatening and non-threatening situations. To safely navigate through the environment, sometimes even to secure survival, we have to identify, memorize, and react to certain cues which indicate potential threats or serve as safety signals. The acquisition, maintenance, and extinction of such conditioned fear responses represent critical cognitive functions, especially in a clinical context with individuals suffering from anxiety disorders (Craske, Hermans, & Vansteenwegen, 2006; Mineka & Oehlberg, 2008), and have been subject to scientific investigation for more than a century. The most commonly used paradigm to study the neural mechanisms underlying fear acquisition and extinction is Pavlovian fear conditioning. Here, subjects are presented with two neutral stimuli, e.g. pictures of differently colored geometric shapes, which are usually referred to as conditioned stimuli (CS). Based on a certain reinforcement rate, one of these stimuli (CS+) is paired with an aversive stimulus, also called the unconditioned or unconditional stimulus (US), while the other conditioned stimulus (CS-) is never paired with the US. As a consequence, subjects learn that the CS+ is indicative of the US and show a conditioned fear response (CR) when the CS+ is presented, but not when the CS- is presented. The CR is usually quantified by measures of skin conductance response (SCR) or subjective fear ratings (Lonsdorf & Merz, 2017).

The combination of Pavlovian fear conditioning and neuroimaging methods such as functional magnetic resonance imaging (fMRI) provides an opportunity to identify specific brain areas involved in fear learning. In this regard, the amygdala became a center piece of the fear conditioning circuit (Milad & Quirk, 2012). However, it has been shown that fear learning involves a whole network of brain regions, which tend to exhibit greater functional activation in response to CS+ presentations compared to CS- presentations. This so called “fear network” is thought to be crucial for identifying and encoding potential threats. According to a recent meta-analysis, it comprises the anterior insular cortex (AIC) extending to the frontal operculum, the ventral striatum and major thalamic nuclei, pre-supplementary and supplementary motor areas, the dorsal anterior cingulate cortex (dACC) and a distinct cluster of the dorsal-anterior precuneus, the second somatosensory cortex, the dorsolateral prefrontal cortex, the lateral premotor cortex, the ventral-posterior precuneus, and the lateral cerebellum (Fullana et al., 2016). Importantly, the authors also described a second brain network comprising brain regions that tend to exhibit greater functional activation in response to CS- presentations compared to CS+ presentations. Brain areas constituting this so called “safety network” are the primary somato-sensory cortex and the dorsal posterior insular cortex, the dorsal anterior prefrontal cortex, the ventromedial prefrontal cortex (vmPFC), the posterior cingulate cortex (PCC) including the retrosplenial cortex, the hippocampus, the lateral orbitofrontal cortex (OFC), the inferior parietal cortex, the posterior cerebellum, the dorsal caudate nucleus, and the dorsal-posterior precuneus. Noteworthy, Fullana et al. (2016) argued that processing non-threatening stimuli that are indicative of safety is not a passive task but requires emotional and cognitive functions. Thus, they considered the safety network to be as important for fear conditioning as the fear network.

Robust models of fear learning can offer valuable insights, especially in a clinical context where they provide a better understanding of the development and progression of anxiety and panic disorders. Unfortunately, the vast majority of these models assumes an idealized “average” individual, while neglecting important mechanisms that lead to interindividual differences in fear response. However, it has been demonstrated that these differences represent stable trait-like qualities (Hartley, Fischl, & Phelps, 2011). In addition, behavioral variations in fear learning have been associated with variations in brain function. For example, Phelps (2004) could show that amygdala activity during fear acquisition is positively correlated to CR, quantified as the difference between mean SCR of CS+ trials and mean SCR of CS- trials (differential CR).

Since it is conceivable that brain function is linked to brain morphology, recent studies have made an effort to identify specific macrostructural brain properties underlying individual fear response. For example, Hartley et al. (2011) could show that cortical thickness of the posterior insula can predict differential CR in a classical fear conditioning paradigm. In another study, cortical thickness of the dACC was shown to be positively associated with fear learning (Milad, Quirk, et al., 2007). However, it is important to note that Milad, Quirk, et al. (2007) did not employ differential CR in order to quantify the participants’ fear response but merely used mean SCR in reaction to CS+ presentations. Following a similar approach, a recent study investigating fear learning in anxiety patients and healthy controls found cortical thickness of the dorsomedial and dorsolateral PFC to be negatively associated with mean SCR in reaction to CS+ and CS- trials (Abend et al., 2020). However, this finding is difficult to interpret, not only because authors refrained from using differential CR as their dependent variable, but also because respective analyses included data from both anxiety patients and healthy controls. With regard to subcortical structures, Cacciaglia, Pohlack, Flor, and Nees (2015) demonstrated that left amygdala volume positively predicts differential CR. In contrast, bilateral hippocampal volumes were shown to be associated with the magnitude of differential contingency ratings, i.e. how well participants could remember which CS was paired with the US and which was not. Furthermore, Cacciaglia et al. (2015) also found a negative association between hippocampal volumes and CS- contingency ratings, meaning that larger hippocampi were related to a more successful identification of safety signals. This result is in line with the observation that the hippocampus is part of a larger brain network processing safety signals (Fullana et al., 2016). Based on their findings, Cacciaglia et al. (2015) argued that autonomic fear response and conscious contingency rating represent two parallel but distinct processes that should be addressed separately. Hippocampal and amygdalar volumes have also been shown to be positively related to mean SCR in reaction to CS+ presentations (Pohlack et al., 2012; Winkelmann et al., 2016). However, Hartley et al. (2011) reported a trend towards a negative association between amygdalar volume and differential CR, although not significant. To summarize, measures of cortical thickness and subcortical brain volume, mainly obtained from fear and safety network components, have already been shown to account for interindividual differences in fear response. However, comparability of respective studies is limited given that some of them related said brain properties to differential CR and others used mean SCR in reaction to CS+ or CS- presentations instead. Moreover, there are mixed results with regard to the neural correlates underpinning objective SCR measures and subjective contingency ratings.

Unfortunately, several of the aforementioned studies on the structural correlates of fear learning followed a region of interest (ROI) based approach with a very limited number of brain areas (Cacciaglia et al., 2015; Hartley et al., 2011; Pohlack et al., 2012). Thereby, they forfeited the possibility to reveal any potential structure-function relationships in other brain areas from the fear and safety networks. In this regard, a more comprehensive approach is to completely refrain from a priori defined ROIs. Instead, brain areas relevant for the analysis of neural correlates potentially underpinning fear learning can be selected based on their functional activation during fear acquisition training. This method can produce different templates, e.g. one for CS+ and one for CS- presentations, which reliably indicate all voxels that exhibit an increased BOLD response during the processing of threat or safety signals respectively. These templates can in turn be used to guide the extraction of mean values from other brain maps representing various neural correlates.

One example of such neural correlates is concentration of gray matter or gray matter density, a measure provided by a method known as voxel-based morphometry (VBM) (Ashburner & Friston, 2000). VBM allows for an analysis of macrostructural differences in brain morphology, i.e. brain volumes, by employing a spatial normalization procedure that registers the T1-weighted brain scans from a group of individuals to a single template. Based on the degree of transformation necessary to fit an individual scan to the template, the value of each voxel, i.e. its gray matter density, is modulated so that no morphometric information from the original scan is lost. Spatially normalized brain scans can be subjected to a voxel-wise analysis, in which gray matter density is related to other (behavioral) variables such as differential CR. Surprisingly, this approach has not been used in existing studies focused on the neural foundations of fear learning so far.

Another realm that has not yet been touched by previous fear learning research is that of microstructural correlates. Among other aspects, this encompasses the morphology of neurites, namely dendrites and axons. In the past, it was not possible to study such properties in vivo due to a lack of suitable brain imaging techniques. Fortunately, technological development during the past decade has spawned several methods capable of providing information about the brain’s architecture at the level of axons or dendrites. Currently, one of the most promising techniques for the quantification of neurite morphology is a diffusion MRI technique known as neurite orientation dispersion and density imaging (NODDI) (Zhang, Schneider, Wheeler-Kingshott, & Alexander, 2012). This technique is based on a multi-shell high-angular-resolution diffusion imaging protocol and offers a novel way to analyze diffusion-weighted data with regard to tissue microstructure. It features a three-compartment model distinguishing intra-neurite, extra-neurite and cerebrospinal fluid (CSF) environments. NODDI is based on a diffusion model that was successfully validated by histological examinations utilizing staining methods in gray and white matter of rats and ferrets (Jespersen et al., 2010; Jespersen, Leigland, Cornea, & Kroenke, 2012). In addition, Zhang et al. (2012) have shown that NODDI is also capable of estimating diffusion markers of neurite density and orientation dispersion by in vivo measurements in humans. Direct validation of NODDI has recently been performed in a study investigating neurite dispersion as a potential marker of multiple sclerosis pathology in post-mortem spinal cord specimens (Grussu et al., 2017). Here, the authors reported a close match between NODDI-derived neurite dispersion and its histological counterpart. This indicates that NODDI metrics are closely reflecting their histological conditions. Moreover, current studies have shown that NODDI predicts interindividual differences in cognitive ability (Genc et al., 2018), language processing (Ocklenburg et al., 2018) and hemispheric asymmetries (Schmitz et al., 2019).

Here, we present the first study to investigate the microstructural underpinnings of fear learning by means of NODDI. In addition, our analyses also employed gray matter density as a potential predictor that has not been related to fear learning in previous research. In order to ensure a comprehensive but precise coverage of both fear and safety network components, we used fMRI task data to select brain regions which exhibited increased functional activation during CS+ or CS- presentations, respectively. We demonstrate that mean gray matter density averaged across brain regions from the fear network is significantly correlated with differential CR as measured by SCR. In contrast, we show that NODDI-derived measures of neurite density and neurite orientation dispersion are not associated with differential CR.

## 2. Material and Methods

### 2.1. Participants

Seventy-six young and healthy participants took part in the current study. In total, sixteen participants had to be excluded due to various reasons. Two participants were removed because of technical difficulties with the electrical stimulation. Three participants were classified as “non-responders” since they did not show valid SCRs in reaction to any of the US presentations (at least 0.05 µS) (Lonsdorf et al., 2019). Six participants reported a CS-/US contingency that was equal to or larger than the reported CS+/US contingency, indicating absent contingency awareness (Tabbert et al., 2011). Four participants were excluded since data on their SCRs to either CS+ or CS- were missing. Lastly, one participant had to be removed due to a corrupted data file containing the diffusion-weighted imaging. Hence, all imaging analyses were carried out with data from the remaining 60 participants (37 women). The age range was 18 to 25 years (*M* = 21.70, *SD* = 1.81). We did not observe significant age differences between male and female participants (*t*(58) = −0.453, *p* = .653).

In order to control for any potential effects caused by handedness, only right-handed individuals were recruited, as measured by the Edinburgh Handedness Inventory (Oldfield, 1971). All participants had normal or corrected-to-normal vision and were able to understand the instructions given to them orally and in writing. They were either paid for their participation or received course credit. All participants were naive to the purpose of the study and had no former experience with the aversive learning paradigm used for the experiment. Participants reported no history of psychiatric or neurological disorders and matched the standard inclusion criteria for fMRI examinations. The study was approved by the local ethics committee of the Faculty of Psychology at Ruhr University Bochum. All participants provided written informed consent prior to participation and were treated in accordance with the Declaration of Helsinki.

### 2.2. Fear Acquisition Paradigm

While in the fMRI scanner, all participants completed differential fear acquisition training followed by fear extinction training. However, as this study is only concerned with fear acquisition, all procedures related to fear extinction will be reported elsewhere. The stimuli and procedure used for fear acquisition training were modified from (Milad, Wright, et al., 2007).

After arrival, participants gave written informed consent and filled out a questionnaire on demographic variables. Prior to scanning, all participants were instructed to pay close attention to the images being presented during fear acquisition training. They were also told that electrical stimulation may or may not be presented during the experiment. Electrical stimulation (1 ms pulses with 50 Hz for a duration of 100 ms) was applied as the US using a constant voltage stimulator (STM2000 BIOPAC systems, CA, USA) along with two electrodes attached to the fingertips of the first and second fingers of the right hand. The intensity of electrical stimulation was adjusted for each participant individually prior to scanning. For this purpose, electrical stimulation was administered at 30 V and raised in increments of 5 V until participants rated the sensation as very unpleasant but not painful.

During fear acquisition training, a picture of an office room with a switched off desk lamp was used as the context image (Figure 1, top right). In each trial, the desk lamp would light up either in blue (CS+) or in yellow (CS-), which served as the two CS. The images were presented using the Presentation software package (Neurobehavioral Systems, Albany, CA) and MR suitable LCD-goggles (Visuastim Digital, Resonance Technology Inc, Northridge, CA). At the beginning of each trial, a white fixation cross was presented on a black background for 6.8 - 9.5 seconds (randomly jittered). Next, the context image was presented for 1 second, which was followed by the presentation of the CS+ or the CS- for another 6 seconds. In the experimental group, the CS+ was paired with electrical stimulation in 62.5% of trials. The stimulation was administered 5.9 seconds after CS+ onset and co-terminated with CS+ offset. Across all 32 trials, CS+ and CS- were presented 16 times each in pseudo-randomized order. The first two trials always consisted of one CS+ and one CS- presentation and so did the last two trials. The first and last CS+ presentations were always paired with electrical stimulation. There were no trials that presented the same type of CS more than twice in consecutive order. CS+ and CS- presentations were distributed equally across both halves of fear acquisition training.

**Figure 1.**
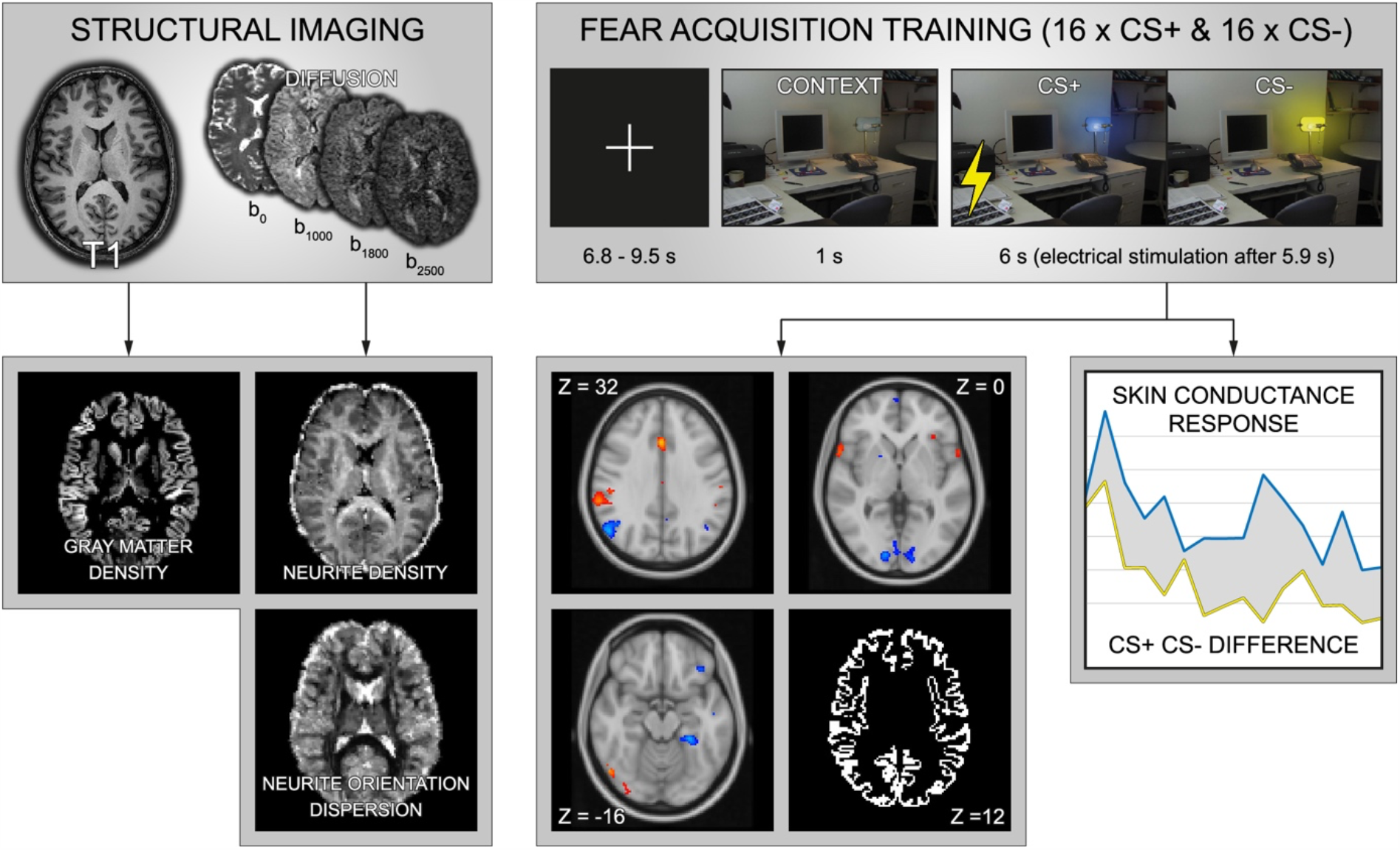
Data acquisition and analysis. All participants had to undergo a structural imaging session, constituted of T1-weighted and diffusion-weighted scans, as well as a fear acquisition training that was conducted during functional magnetic resonance imaging. T1-weighted images were used to compute brain maps of gray matter density. Diffusion-weighted images were obtained using multiple shells (b = 0 mm/s^2^, b = 1000 mm/s^2^, b = 1800 mm/s^2^, b = 2500 mm/s^2^) and subjected to an analysis of neurite density and neurite orientation dispersion. Fear acquisition training comprised 32 trials in total. At the beginning of each trial, a white fixation cross was presented on a black background for 6.8 - 9.5 seconds. Next, the context image was presented for 1 second, which was followed by the presentation of the CS+ (blue light) or the CS- (yellow light) for another 6 seconds. The CS+ was paired with electrical stimulation in 62.5% of trials. The stimulation was administered 5.9 seconds after CS+ onset. The CS+ and CS- were presented 16 times each in pseudo-randomized order. The degree of conditioned reaction was quantified by subtracting average skin conductance responses to the CS- from average responses to the CS+. Voxel clusters exhibiting statistically increased BOLD responses in reaction to CS+ and CS- trials were used to generate two separate brain masks. Respective “CS+ > CS-” and “CS- > CS+” masks were restricted to the cortical ribbon. For each participant, mean values of gray matter density, neurite density, and neurite orientation were computed by averaging across all voxels from the “CS+ > CS-” and “CS- > CS+” masks, respectively. These mean values were correlated with conditioned reaction measures while controlling for confounding effects of age and sex.

After fear acquisition training, participants had to rate the contingencies between CS+, CS-, and US. First, they were asked to report the number of electrical stimulations they had received during the experiment. In addition, they were asked to rate the unpleasantness of the last electrical stimulation on a 9-point Likert scale (1 - “not unpleasant”, 9 - “very unpleasant”) and to report at what rate the blue lamplight and the yellow lamplight had been followed by an electrical stimulation.

### 2.3. Acquisition of Imaging Data

All imaging data were acquired at the Bergmannsheil hospital in Bochum, Germany, using a 3T Philips Achieva scanner with a 32-channel head coil. Scanning included T1-weighted imaging, diffusion-weighted imaging (multiple shells), and task-based imaging (fear acquisition training). All images were obtained during one consecutive session with structural imaging (T1-weighted and diffusion-weighted imaging) preceding task-based imaging during fear acquisition training (Figure 1, top).

#### 2.3.1. T1-weighted Imaging

For the purpose of coregistration and for computing gray matter density, T1-weighted high-resolution anatomical images were acquired (MP-RAGE, TR = 8.2 ms, TE = 3.7 ms, flip angle = 8°, 220 slices, matrix size = 240 × 240, voxel size = 1 mm × 1 mm × 1 mm). Scanning time was around 6 minutes (Figure 1, top left).

#### 2.3.2. Diffusion-weighted Imaging

For the analysis of NODDI coefficients, diffusion-weighted images were acquired using echo planar imaging (TR = 7652 ms, TE = 87 ms, flip angle = 90°, 60 slices, matrix size = 112 × 112, voxel size = 2 mm × 2 mm × 2 mm). Diffusion weighting was based on a multi-shell, high angular resolution scheme consisting of diffusion-weighted images for b-values of 1000, 1800, and 2500 s/mm^2^, respectively, applied along 20, 40, and 60 uniformly distributed directions. All diffusion directions within and between shells were generated orthogonal to each other using the MASSIVE toolbox (Froeling, Tax, Vos, Luijten, & Leemans, 2016). Additionally, eight volumes with no diffusion weighting (b = 0 s/mm^2^) were acquired as an anatomical reference for motion correction and computation of NODDI coefficients. Diffusion-weighted data was collected with reversed phase-encode directions, resulting in pairs of images with distortions going in opposite directions. The acquisition time of the diffusion-weighted images was 18 minutes (Figure 1, top left).

#### 2.3.3. Task-based Imaging

In order to identify specific brain regions showing a significantly pronounced or diminished BOLD response during fear acquisition, we employed echo planar imaging. We obtained a time series of fMRI volumes for each participant while completing fear acquisition training (TR = 2500 ms, TE = 35 ms, flip angle = 90°, 40 slices, matrix size = 112 × 112, voxel size = 2 mm × 2 mm × 3 mm). Scanning time was around 8 minutes.

### 2.4. Acquisition of Skin Conductance Responses

Skin conductance responses were assessed using two Ag/AgCl electrodes filled with isotonic (0.05 NaCl) electrolyte medium placed on the hypothenar eminence right below the fifth finger of the left hand. Data were recorded using Brain Vision Recorder software (Brain Products GmbH, Munich, Germany).

### 2.5. Preprocessing of Imaging Data

#### 2.5.1. Preprocessing of T1-weighted Data

In order to create maps of gray matter density based on T1-weighted data, we utilized FSL-VBM (Douaud et al., 2007), which is an optimized voxel-based morphometry protocol (Good et al., 2001) carried out with FSL tools (Smith et al., 2004). First, structural images were brain-extracted and gray matter-segmented before being registered to the MNI 152 standard space using non-linear registration (Andersson, Jenkinson, & Smith, 2007). The resulting images were averaged and flipped along the x-axis to create a left-right symmetric, study-specific gray matter template. Second, all native gray matter images were non-linearly registered to this study-specific template and “modulated” to correct for local expansion (or contraction) due to the non-linear component of the spatial transformation. Finally, the modulated gray matter images were non-linearly registered to the MNI 152 standard space template.

#### 2.5.2. Preprocessing of Diffusion-weighted Data

Diffusion-weighted data were prepared for the computation of NODDI coefficients by means of a preprocessing pipeline with following steps. First, images were corrected for signal drift (Vos et al., 2017) using ExploreDTI (Leemans, Jeurissen, Sijbers, & Jones, 2009). Second, we utilized the topup tool as implemented in FSL (Smith et al., 2004), a method similar to that described in (Andersson, Skare, & Ashburner, 2003), to estimate the susceptibility-induced off-resonance field based on pairs of images with opposite phase-encode directions (2.3.3. Diffusion-weighted Imaging). Third, the topup output was used in combination with eddy (Andersson & Sotiropoulos, 2016), which is also part of the FSL toolbox, to correct for susceptibility, eddy currents, and subject movement. Importantly, we also performed outlier detection as implemented in eddy to identify slices where signal had been lost as a consequence of subject movement during the diffusion encoding (Andersson, Graham, Zsoldos, & Sotiropoulos, 2016).

NODDI coefficients were computed from preprocessed diffusion-weighted data using the AMICO toolbox (Daducci et al., 2015). The AMICO approach is based on a convex optimization procedure that converts the non-linear fitting into a linear optimization problem (Daducci et al., 2015). This reduces processing time dramatically (Sepehrband, Alexander, Kurniawan, Reutens, & Yang, 2016). The NODDI technique features a three-compartment model distinguishing intra-neurite, extra-neurite and CSF environments. For this study, we decided to focus on the intra-neurite compartment exclusively. Intra-neurite environments are characterized by a stick-like or cylindrically symmetric diffusion signal that is created when water molecules are restricted by the membranes of neurites. In white matter this kind of diffusion is likely to emerge from the presence of axons. In gray matter it serves as an indicator of dendrites and axons forming the neuropil. For each voxel, NODDI computes the intra-neurite volume fraction (INVF), i.e. the portion of the diffusion signal that can be attributed to intra-neurite environments (Jespersen et al., 2010; Jespersen et al., 2012; Zhang et al., 2012). This measure is used to quantify neurite density. In addition, NODDI provides a tortuosity measure coupling the intra-neurite space and the extra-neurite space. This metric, known as the neurite orientation dispersion index (ODI), is supposed to reflect the alignment or dispersion of axons in white matter or axons and dendrites in gray matter (Billiet et al., 2015; Zhang et al., 2012). Examples of INVF and ODI maps from a representative individual are illustrated in Figure 1 (bottom left). Similar to gray matter density maps, the INVF and ODI maps of each individual were non-linearly registered to the MNI 152 standard space template.

#### 2.5.3. Preprocessing of Task-based Data

Task-based data were analyzed by means of FEAT from the FSL toolbox (Smith et al., 2004). Pre-processing of respective images involved motion- and slice-timing correction, spatial smoothing with a 6 mm FWHM Gaussian kernel, high-pass filtering with the cutoff set to 50 seconds, linear registration to the individual’s high-resolution T1-weighted anatomical image, and non-linear registration to the MNI 152 standard space template. We performed first-level analyses (Woolrich, Ripley, Brady, & Smith, 2001) to generate two statistical maps of functional activation (CS+ > CS- and CS- > CS+) including data from all 16 CS+ and 16 CS- presentations. All regressors (context alone, CS+, CS-, US, US omission after CS+ presentation, non-US after CS- presentation) were included in a general linear model based on a stick function that was convolved with the canonical hemodynamic response function, without specifically modeling the durations of the different events (i.e. event-related design). Second-level analyses (Woolrich, Behrens, Beckmann, Jenkinson, & Smith, 2004) employed random-effects estimation by means of FLAME (FMRIB’s Local Analysis of Mixed Effects). The primary effects of interest included both of the aforementioned contrasts (CS+ > CS- and CS- > CS+) for the whole brain volume. For these contrasts we utilized an FWE-corrected cluster thresholding option, *p*-values < .05, and *Z*-values > 3.1.

#### 2.3.4. Creation of Cortical Brain Masks

For the purpose of masking aforementioned brain maps of gray matter density, neurite density, and neurite orientation dispersion, we utilized the standard cortical surface in FreeSurfer called fsaverage. The fsaverage parcellation, covering the cortical ribbon, was registered to the MNI 152 standard space template and converted to a volumetric mask. Each of the two brain maps reflecting the contrasts in functional activation between CS+ and CS- presentations (see 2.5.3. Preprocessing of Task-based Data) were binarized and multiplied with the aforementioned cortical mask. This resulted in two brain masks covering cortical areas in which increases of the BOLD signal in response to either CS+ or CS- were observed during fear acquisition training.

### 2.6. Analysis of Skin Conductance Responses

Raw SCR data were pre-processed with Brain Vision Analyzer software (Brain Products GmbH, Munich, Germany). All further analyses were conducted semi-automatically using MATLAB, version 9.6.0.1114505 (R2019a, The MathWorks Inc., Natrick, MA). SCR to CS presentation was defined as the maximum amplitude recorded within the time window starting 1 s after CS onset and ending 6.5 s after CS onset. The CR was quantified as the difference between the average SCR across CS+ trials and the average SCR across CS- trials (Figure 1, bottom right). SCR to US presentation was defined as the maximum amplitude recorded within the time window starting 6.5 s after CS onset and ending 12 s after CS onset.

### 2.7. Statistical Analysis

All statistical analyses were carried out using MATLAB, version 9.3.0.713579 (R2017b, The MathWorks Inc., Natick, MA). For all analyses, we employed linear parametric methods. Testing was two-tailed with an α-level of .05, which was corrected for multiple comparisons. The current study aimed to investigate the association between structural brain markers and fear learning in functionally defined regions of interest involved in the processing of threatening and non-threatening stimuli. To this end we utilized the two brain masks that were created based on the functional contrasts which resulted from fear acquisition training (2.3.4. Creation of Cortical Brain Masks). The gray matter density, INVF, and ODI maps of each participant were multiplied with the fear (CS+ > CS-) and safety (CS- > CS+) network masks, respectively. The remaining voxels were averaged to create six mean coefficients (GMD_fear_, GMD_safety_, INVF_fear_, INVF_safety_, ODI_fear_, ODI_safety_).

First, in order to check for multicollinearity among the variables, we correlated all of the aforementioned structural coefficients with each other. At this stage, we did not correct α-levels for multiple comparisons since we also aimed to detect weak associations between variables, even those failing statistical significance by a narrow margin.

Second, we analyzed the relationship between the structural coefficients and conditioned fear reaction by computing Pearson correlations and controlling for potentially confounding effects of age and sex. Here, the α-levels applied to the resulting partial correlations were corrected using the Bonferroni method (α* = .05 / 6 = .0083). Third, we performed multiple regression analysis in order to account for any correlations among the structural coefficients potentially distorting the genuine contribution of each coefficient towards conditioned fear reaction. To this end, we created a regression model with conditioned fear reaction as the dependent variable and all six structural coefficients as well as age and sex serving as independent variables.

## 3. Results

### 3.1. Fear Memory Acquisition

Participants showed significantly greater SCRs in reaction to CS+ presentations relative to CS- presentations (*t*(59) = 6.015, *p* < .001) (Figure 2), indicating successful fear learning.

**Figure 2.**
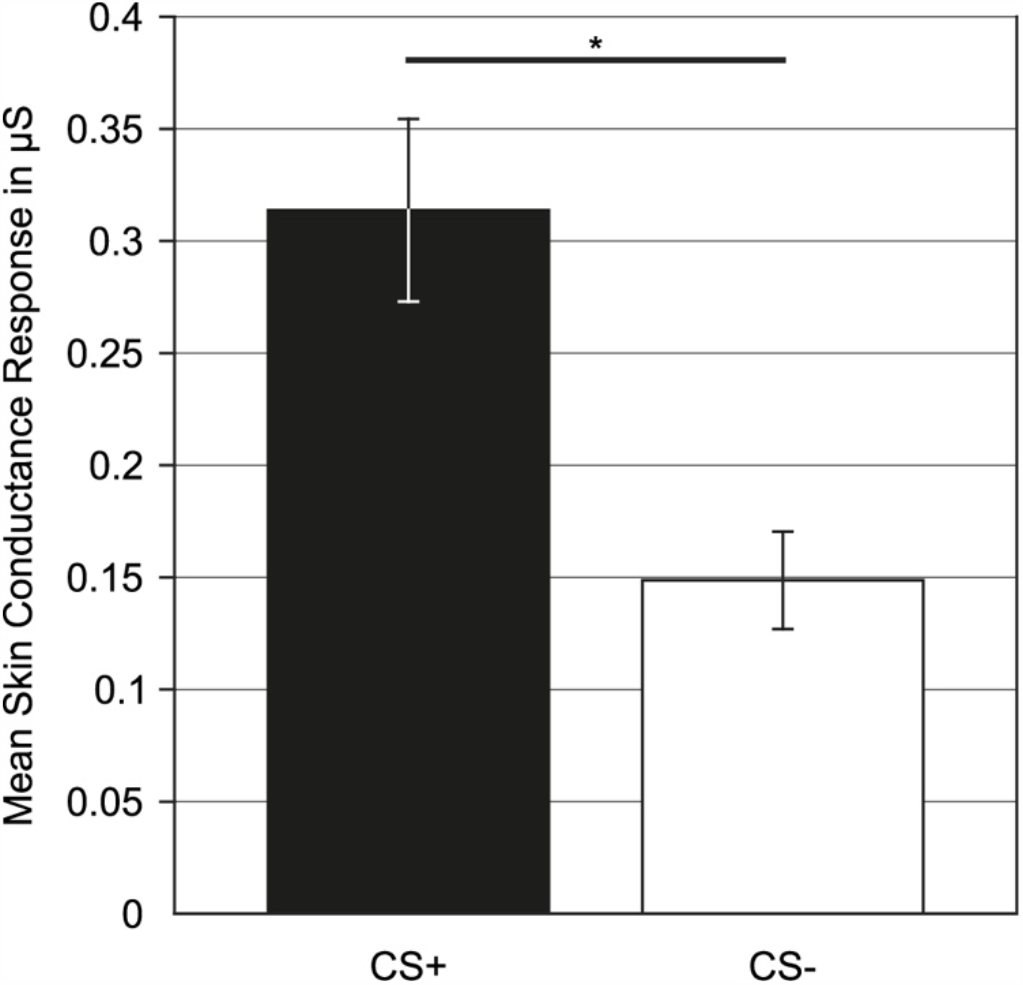
Mean skin conductance responses. Bar plots showing the mean skin conductance responses in reaction to CS+ (black bar) and CS- (white bar) presentations in microSiemens (μS). Data were averaged across all 16 CS+ and 16 CS- trials, respectively. Error bars represent standard errors (* p < .001).

Furthermore, fMRI data obtained during fear acquisition training revealed distinct patterns of functional activation in reaction to CS+ and CS- presentations. Overall, our data were in good accordance with the results of a large-scale meta-analysis on the functional correlates of fear acquisition (Fullana et al., 2016). Voxel clusters exhibiting significant (*p* < .05, *Z* > 3.1, FWE-corrected) BOLD signal contrasts in our data either overlapped with components of the so-called fear (CS+ > CS-) and safety (CS- > CS+) networks or were directly adjacent to them (Supplementary Figure 1 and Supplementary Figure 2). With regard to the fear network, we were able to replicate relevant structures like the AIC (frontal operculum), dACC, thalamus, and second somatosensory cortex (functional activation in the CS+ > CS- contrast). Likewise, relevant structures from the safety network, namely the inferior parietal cortex, primary somatosensory cortex, parahippocampal gyrus, PCC, vmPFC, and lateral OFC, could also be identified in our data (functional activation in the CS- > CS+ contrast).

### 3.2. Partial Correlation Analysis

We used aforementioned fMRI data showing functional activation in reaction to CS+ or CS- presentations to create templates for the extraction of mean values from gray matter density, neurite density, and neurite orientation dispersion maps. Given that functionally active voxel clusters observed in our data were in good accordance with the fear and safety networks described by Fullana et al. (2016), respective mean values were named GMD_fear_, GMD_safety_, INVF_fear_, INVF_safety_, ODI_fear_, and ODI_safety_. Previous research has demonstrated that age and sex are significantly associated with measures of brain volume (Genc et al., 2019) and NODDI coefficients (Genc et al., 2018). Hence, we decided to control for potentially confounding effects of age and sex by employing partial correlations. In total, we computed six correlations between the structural coefficients and conditioned fear reaction with age and sex being partialled out. Neither INVF_fear_ (r = -.034, p = .802) and INVF_safety_ (r = -.110, p = .413) nor ODI_fear_ (r = .103, p = .442) and ODI_safety_ (r = -.080, p = .550) exhibited statistically significant associations with conditioned fear response. In view of gray matter density, we observed conditioned fear reaction to be significantly correlated with GMD_fear_ (r = .361, p = .007) (Figure 3) but not GMD_safety_ (r = -.183, p = .170).

**Figure 3.**
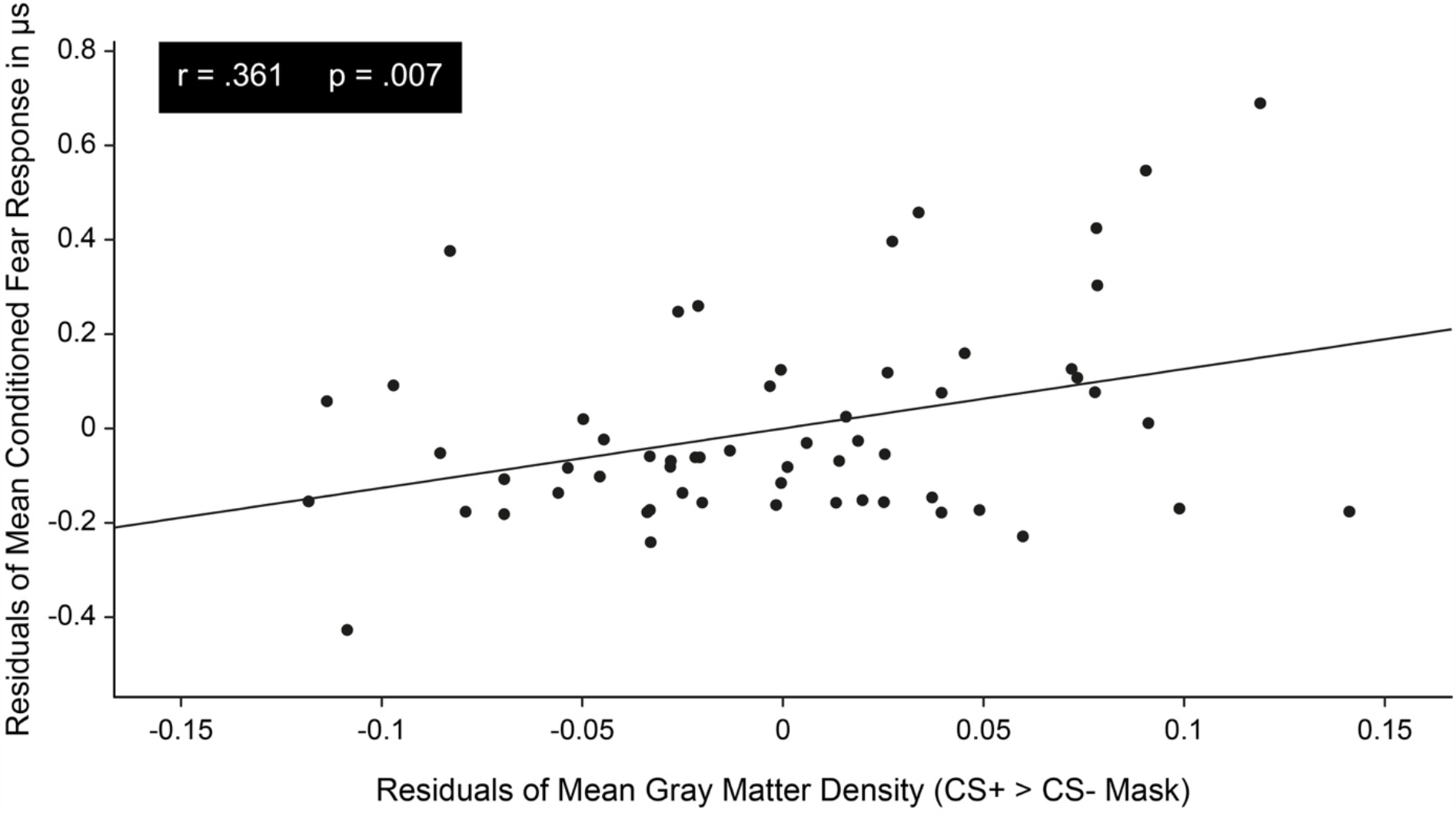
Partial regression diagram showing the relationship between gray matter density (“CS+ > CS-” mask) and mean conditioned fear response. The scatter plot illustrates the result yielded by a partial correlation analysis controlling for potential confounding effects of age and sex. Mean gray matter density coefficients were computed by averaging the values of all voxels from the gray matter density map corresponding to the “CS+ > CS-” mask, i.e. voxels exhibiting a significantly higher BOLD response in reaction to CS+ presentations compared to CS- presentations. Mean conditioned fear responses were computed by subtracting the average skin conductance responses across all CS- trials from the average skin conductance responses across all CS+ trials.

### 3.3. Multiple Regression Analysis

In order to check our data for multicollinearity, we computed Pearson correlations for all possible combinations between the six structural coefficients (Supplementary Table 1). Five of these correlations turned out to be statistically significant. GMD_fear_ was significantly correlated with INVF_fear_ (r = -.327, p < .05) and INVF_safety_ (r = -.262, p < .05), indicating a general inverse association regardless of neurite density being obtained from the CS+ > CS- or CS- > CS+ mask. In addition, GMD_fear_ also exhibited a significant positive correlation with ODI_safety_ (r = .307, p < .05). We observed a highly significant positive correlation between INVF_fear_ and INVF_safety_ (r = .552, p < .001), indicating similar degrees of neurite density in brain regions involved in the processing of threat and safety signals. However, we found ODI_safety_ to be significantly correlated with INVF_fear_ (r = -.323, p < .05) but not INVF_safety_. It is noteworthy that almost all of the correlations initially reaching statistical significance did not survive a Bonferroni correction for multiple comparisons (α* = .05 / 28 = .0019). The only exception was the association between the INVF_fear_ and INVF_safety_ coefficients. Nevertheless, we decided to treat these associations as potential confounding factors and addressed this issue by computing a multiple regression analysis. Conditioned fear reaction served as the dependent variable and was regressed on all six structural coefficients as well as age and sex (Figure 4). This regression model confirmed the results obtained from our partial correlation analysis. GMD_fear_ turned out to be the only significant predictor of conditioned fear reaction (*β* = .436, p < .01), whereas the associations between conditioned fear reaction and all other structural coefficients as well as age and sex did not reach statistical significance. Overall, this regression model was able to explain 22.60% of variance in conditioned fear reaction.

**Figure 4.**
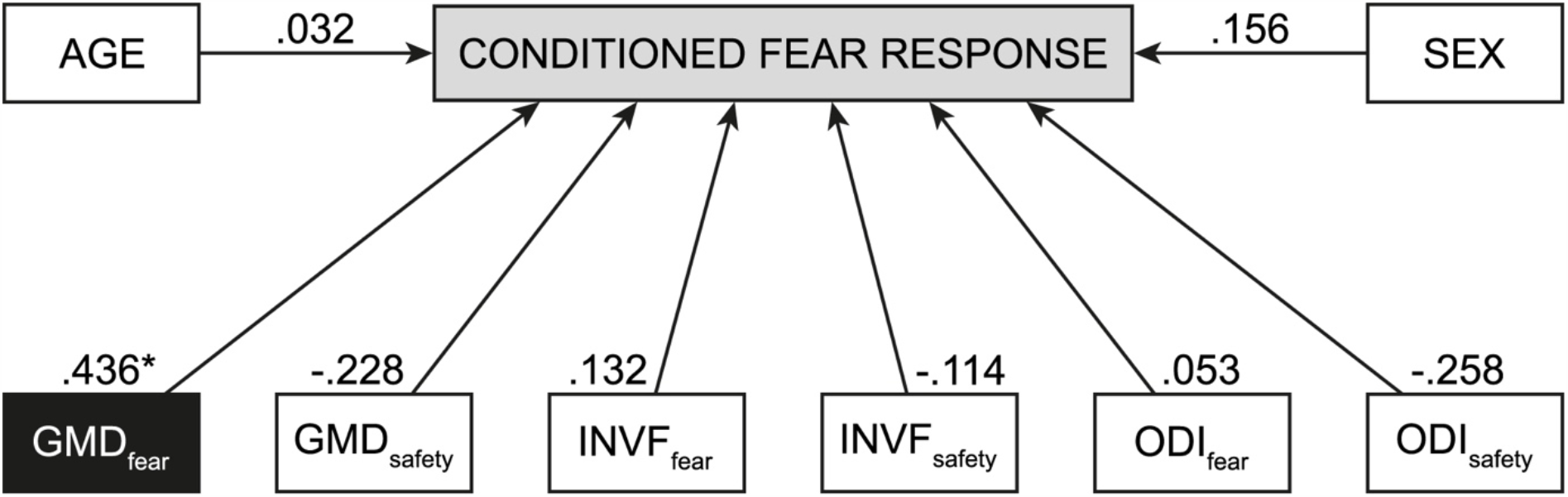
Results of multiple regression analysis. Conditioned fear response as measured via skin conductance responses was regressed on age, sex, and six structural brain properties (GMD_fear_ = gray matter density averaged across the fear network, GMD_safety_ = gray matter density averaged across the safety network, INVF_fear_ = neurite density averaged across the fear network, INVF_safety_ = neurite density averaged across the safety network, ODI_fear_ = neurite orientation dispersion averaged across the fear network, ODI_safety_ = neurite orientation dispersion averaged across the safety network). Standardized regression coefficients are depicted next to the respective arrows. * p < .01.

## 4. Discussion

The goal of this study was to extend our knowledge about the macrostructural underpinnings of fear learning as well as to gain new insights regarding potential microstructural correlates. In contrast to previous research, we did not select regions of interest based on a priori assumptions. Instead, we conducted fear acquisition training while recording fMRI data. By so doing, we were able to identify clearly defined voxel clusters showing increased functional activation during CS+ or CS- presentations, i.e. the fear and safety networks. For our analyses we utilized respective network templates to obtain mean values of different structural coefficients, which had not been associated with fear learning in previous research so far. On the one hand, we employed gray matter density as a macrostructural measure of brain morphometry. On the other hand, we used two NODDI coefficients, supposed to capture cortical microstructure in the form of neurite density and neurite orientation dispersion. We could show that mean gray matter density averaged across the fear network is positively associated with differential CR, whereas no such relation was observed for mean gray matter density obtained from the safety network. Furthermore, our analysis of NODDI-derived measures revealed no significant associations between fear learning and the microstructural architecture of the cortex.

At its core, gray matter density is an indicator of regional brain volume. Due to the processing pipeline employed by voxel-based morphometry, higher values of gray matter density are likely to represent cortical segments with increased thickness or surface area, especially in healthy individuals without any forms of neurodegeneration. Hence, our observation that differential CR can be predicted by mean gray matter density averaged across the fear network is in line with previous studies on the structural basis of fear learning. Among other regions, our fear network template comprised parts of the AIC and dACC. In previous research cortical thickness of the posterior insula was associated with differential CR (Hartley et al., 2011) and cortical thickness of the dACC was associated with mean SCR in reaction to CS+ presentations (Milad, Quirk, et al., 2007). In general, these results appear plausible given that a larger cortical volume is likely to hold more neurons, which in turn provides more computational power to process relevant information. For example, higher cognitive functions such as mental reasoning have consistently been shown to benefit from larger brain volume (Pietschnig, Penke, Wicherts, Zeiler, & Voracek, 2015). With regard to fear learning, an enlarged fear network might possess an increased cognitive capacity to process threatening stimuli like the CS+. This could lead to a more pronounced fear response in terms of SCR, which automatically precipitates a better differentiation between threatening and non-threatening stimuli, i.e. differential CR.

In contrast to the fear network, the safety network did not exhibit a significant association between gray matter density and differential CR. Our safety network template was based on voxel clusters showing functional activation for the contrast CS- > CS+. Hence, it is surprising that an increase of gray matter density in respective brain regions did not lead to a more efficient processing of safety signals, which would have decreased mean SCR in reaction to CS- presentations and in turn increased differential CR. This lack of an association is also reflected in the fact that the variance of mean SCR during CS- presentations was smaller compared to that during CS+ presentations (Figure 2). During fear acquisition training the CS- was never paired with electric stimulation. Hence, it is conceivable that participants did not show much interindividual variation in their respective SCR because identifying the CS- as a safety signal did not present a very challenging task. In contrast, the partial reinforcement plan for CS+ presentations made it more difficult to categorize the stimulus as a threat or safety signal and might be the reason for a more pronounced variance in respective mean SCR. It may be possible that differential CR was not affected by changes in the safety network’s gray matter density because the task of identifying the CS- as a safety signal did not have the potential to be carried out more efficiently in the first place.

With regard to NODDI-derived measures, we did not find neurite density or neurite orientation dispersion to be significantly related to differential CR. To the best of our knowledge, we conducted the first study to employ NODDI as a method of investigating the microstructural foundations of fear learning. However, previous research from different fields has related neurite density to other cognitive processes. For example, Genc et al. (2018) reported a negative association between neurite density and fluid intelligence, indicating that higher cognitive functions like mental reasoning might benefit from a sparse but well-organized cortical microstructure. In contrast, Ocklenburg et al. (2018) found that comparatively basic cognitive functions like speech processing are facilitated by a higher neurite density in the planum temporale. Based on these findings, it seems likely that different types of cognitive processes benefit from different types of microstructural architecture. Given that fear conditioning can be considered a rather basic cognitive process, we would have expected results similar to those reported by Ocklenburg et al. (2018), namely a positive relationship between differential CR and neurite density in the fear and safety networks. However, the absence of any significant findings might indicate one of two things. First, it is entirely possible that interindividual differences in fear learning are not driven by any variation in cortical microstructure, at least not by measures of neurite density or neurite orientation dispersion. Second, if differences in cortical microstructure do have an impact on fear learning, it could be that our approach was not capable of properly quantifying such differences. With regard to the latter scenario, we still believe that it is preferable to define regions of interest by means of task-based fMRI data instead of a priori assumptions.

Nevertheless, with NODDI-derived measures it could be possible that too much information is lost in the process of averaging data across different brain regions. Although Genc et al. (2018) found fluid intelligence to be significantly associated with mean neurite density averaged across the whole cortex, the microstructural correlates of fear conditioning might follow a more heterogenous distribution. We know that different modules of the fear and safety networks regulate different aspects of fear learning and that their interplay and communication rely on complex excitatory and inhibitory connections (Quirk & Mueller, 2008). Therefore, it is conceivable that some network components might profit from low neurite densities, whereas the opposite might be the case for other brain regions. In such a case, our approach of examining mean neurite density and mean neurite orientation dispersion averaged across whole networks would be prone to disguise potential effects embedded in the microstructure of distinct network components. Hence, for future research we suggest a voxel-wise examination of NODDI-derived measures within regions of interest defined by task-based fMRI data. Furthermore, research on the microstructural correlates underpinning fear learning is likely to profit from imaging data with higher spatial resolution. Recently, it has been shown that ultra-high field MRI with a magnetic field of 7 T is able to provide diffusion-weighted data that can be utilized to examine the microstructural properties of individual cortical layers or subcortical nuclei (Gulban et al., 2018). In the near future, this approach might generate unprecedented insights into the microstructural architecture underlying fear conditioning.

In conclusion, our study was successful in identifying a new structural correlate underlying interindividual differences in fear learning, namely gray matter density. Importantly, these results were not obtained by analyzing data from single a priori defined brain regions but from whole brain networks involved in the processing of threat and safety signals. Moreover, our study is the first to employ neurite imaging for the purpose of investigating potential brain correlates of fear learning but failed to find any significant associations between cortical microstructure and differential CR. However, neurite imaging might benefit from data with higher spatial resolution collected from bigger samples in future research.

## 5. Acknowledgements

This work is part of the SFB 1280 projects A02, A03, A09, and F02 and was supported by the Deutsche Forschungsgemeinschaft (DFG) (project number 31680338). The authors thank M. A. Fullana and B. J. Harrison for providing us with the original maps of functional brain activation from their meta-analysis on human fear conditioning. Further, the authors thank PHILIPS Germany (Burkhard Mädler) for scientific support with the MRI measurements as well as Tobias Otto for technical support. Finally, the authors would like to thank all research assistants for their support with data acquisition.

