## Supplementary Figure 1 for "Conditioned fear reactions are associated with gray matter density but not cortical microstructure"

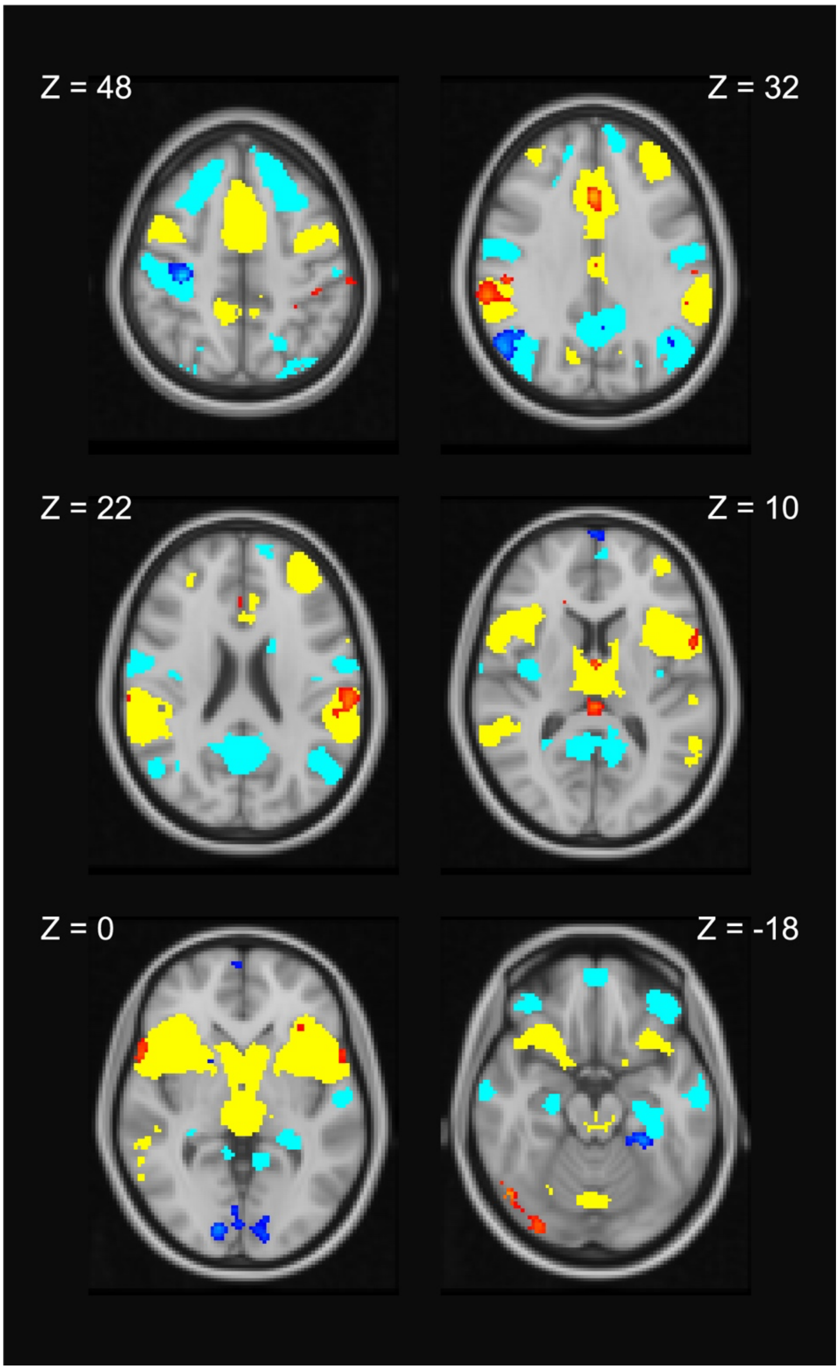


**Supplementary Figure 1. Significant brain activation during fear acquisition training (axial view).** Voxels demonstrating significant BOLD signal contrasts between CS+ and CS- ($p < .05$, $Z > 3.1$, FWE-corrected) are displayed on a selection of axial slices from the Montreal Neurological Institute 152 T1 2-mm template. Brain activation in response to the CS+ versus the CS- is depicted in red to yellow colors, while brain activation in response to the CS- versus the CS+ is depicted in blue to light blue colors. Uniformly colored voxel clusters represent CS+ > CS- activation (yellow) and CS- > CS+ activation (light blue) as reported in a meta-analysis of fMRI studies on fear conditioning (Fullana et al., 2016).
