## Supplementary Table 1 for "Conditioned fear reactions are associated with gray matter density but not cortical microstructure"

| | $GMD_{fear}$ | $GMD_{safety}$ | $INVF_{fear}$ | $INVF_{safety}$ | $ODI_{fear}$ |
|---|---|---|---|---|---|
| **$GMD_{fear}$** | - | - | - | - | - |
| **$GMD_{safety}$** | -.012 | - | - | - | - |
| **$INVF_{fear}$** | -.327* | .122 | - | - | - |
| **$INVF_{safety}$** | -.262* | .109 | .552** | - | - |
| **$ODI_{fear}$** | .078 | -.145 | -.034 | -.015 | - |
| **$ODI_{safety}$** | .307* | -.211 | -.323* | -.212 | .203 |

**Supplementary Table 1. Associations among structural coefficients.** The table shows Pearson correlations between mean structural coefficients obtained from the fear network (CS+ > CS- template) and the safety network (CS- > CS+ template). GMD = gray matter density, INVF = neurite density, ODI = neurite orientation dispersion. * = $p < .05$, ** = $p < .01$
